# Genetic suppression interactions are highly conserved across genetic backgrounds

**DOI:** 10.1101/2024.08.28.610086

**Authors:** Claire Paltenghi, Jolanda van Leeuwen

## Abstract

Genetic suppression occurs when the phenotypic defects caused by a deleterious mutation are rescued by another mutation. Suppression interactions are of particular interest for genetic diseases, as they identify ways to reduce disease severity, thereby potentially highlighting avenues for therapeutic intervention. To what extent suppression interactions are influenced by the genetic background in which they operate remains largely unknown. However, a high degree of suppression conservation would be crucial for developing therapeutic strategies that target suppressors. To gain an understanding of the effect of the genetic context on suppression, we isolated spontaneous suppressor mutations of temperature sensitive alleles of *SEC17, TAO3*, and *GLN1* in three genetically diverse natural isolates of the budding yeast *Saccharomyces cerevisiae*. After identifying and validating the genomic variants responsible for suppression, we introduced the suppressors in all three genetic backgrounds, as well as in a laboratory strain, to assess their specificity. Ten out of eleven tested suppression interactions were conserved in the four yeast strains, although the extent to which a suppressor could rescue the temperature sensitive mutant varied across genetic backgrounds. These results suggest that suppression mechanisms are highly conserved across genetic contexts, a finding that is potentially reassuring for the development of therapeutics that mimic genetic suppressors.

## INTRODUCTION

Predicting phenotype from genotype remains challenging. Although some mutations, such as Mendelian disease alleles, are detrimental in nearly all individuals, the phenotype of most mutations is influenced by their environmental or genetic context, complicating the prediction of a mutation’s phenotype (Nadeau 2001, Chandler et al. 2013, Cooper et al. 2013, Busby et al. 2019, Turco et al. 2023). Genetic context-dependency arises when modifying mutations either increase the severity of a genetic trait or protect against the deleterious effects of a particular mutation (Genin et al. 2008, Harper et al. 2015). Protective modifiers, also called suppressors, can occur in the same gene as the detrimental mutation, or may affect another gene (Lehner 2011, Van Leeuwen et al. 2017). Because suppressors can rescue deleterious phenotypes, suppressors of disease alleles may reveal new therapeutic avenues for treating genetic diseases (Esrick et al. 2021, Frangoul et al. 2021, Ünlü et al. 2023). For example, loss-of-function variants in *BCL11A*, encoding a transcriptional repressor of fetal hemoglobin subunit γ, lead to expression of this subunit in adults. The γ subunit can functionally replace the hemoglobin β subunit, which is compromised in β-thalassemia patients, thereby protecting carriers of *BCL11A* loss-of-function variants against severe β-thalassemia (Uda et al. 2008). This finding led to the development of a gene editing therapy targeting *BCL11A* (Frangoul et al. 2021), which was recently approved for treating β-thalassemia. Despite the success of this therapy aimed at a genetic suppressor, for suppressors to be widely adopted for clinical targeting, they must be conserved across individuals with diverse genetic backgrounds. However, to what extent suppression interactions are influenced by the genetic context in which they operate remains unknown.

Previous studies, focused on the genetic context dependency of particular genetic interactions of interest, have mainly described large differences in interactions between genetic backgrounds (Chari and Dworkin 2013, Wang et al. 2013, Filteau et al. 2015, Mullis et al. 2018). Similarly, a systematic study of the genetic interactions of three yeast genes involved in sterol homeostasis in four genetically diverged yeast strains found that the vast majority of synthetic sick or lethal interactions, in which the combination of two viable mutants leads to a severe fitness defect or lethality, were unique to one genetic background (Busby et al. 2019). However, the generality of these findings for genes involved in other cellular processes remains uncertain. Furthermore, compared to other types of genetic or physical interactions, extragenic suppression interactions are relatively rare and highly enriched for connecting genes that function in the same protein complex or pathway (Van Leeuwen et al. 2016, Van Leeuwen et al. 2020). These properties of genetic suppression may lead to differences in genetic background dependency compared to other types of interactions.

Here, we harnessed the powerful genetics of the budding yeast *Saccharomyces cerevisiae* to study the genetic context-dependency of suppression interactions. We find that the vast majority of identified interactions were conserved in the four tested genetic backgrounds. Nonetheless, the strength of the suppression phenotype varied across contexts, and was sometimes dependent on the sequence or expression level of the suppressor allele. These results suggest that suppression interactions are highly conserved across genetic backgrounds, but that the extent of suppression is influenced by additional genetic variants present in the genome.

## RESULTS

### Systematic identification of genetic suppressors

To study the conservation of suppression interactions across yeast strains, we selected three functionally diverse “query” genes (*SEC17, TAO3*, and *GLN1*). The three query genes are essential for cell viability and are involved in the fusion of vesicles transiting between organelles (*SEC17*) (Clary et al. 1990), regulation of the RAM signaling network for cell proliferation (*TAO3*) (Nelson et al. 2003), and the synthesis of glutamine (*GLN1*) (Mitchell 1985). We used six sequential backcrosses to transfer temperature sensitive (TS) alleles of the three query genes from the S288C reference background into three genetically diverse budding yeast strains of distinct geographical locations and sources: L-1374, UWOPS87-2421, and NCYC110 (**Fig. 1A**) (Liti et al. 2009). The three yeast strains have a nucleotide divergence of 0.40, 0.59, and 0.69%, respectively, compared to the reference strain. After six backcrosses, ∼98% of this genetic divergence should be maintained. All TS alleles still showed a temperature sensitive phenotype in the various strain backgrounds (**Fig. S1**). For *TAO3*, however, the restrictive temperature of the *tao3-5010* allele varied from 30°C in UWOPS87-2421 to 38°C in S288C, suggesting that the severity of the allele was affected by genomic variants present in these strains.

**Figure 1.**
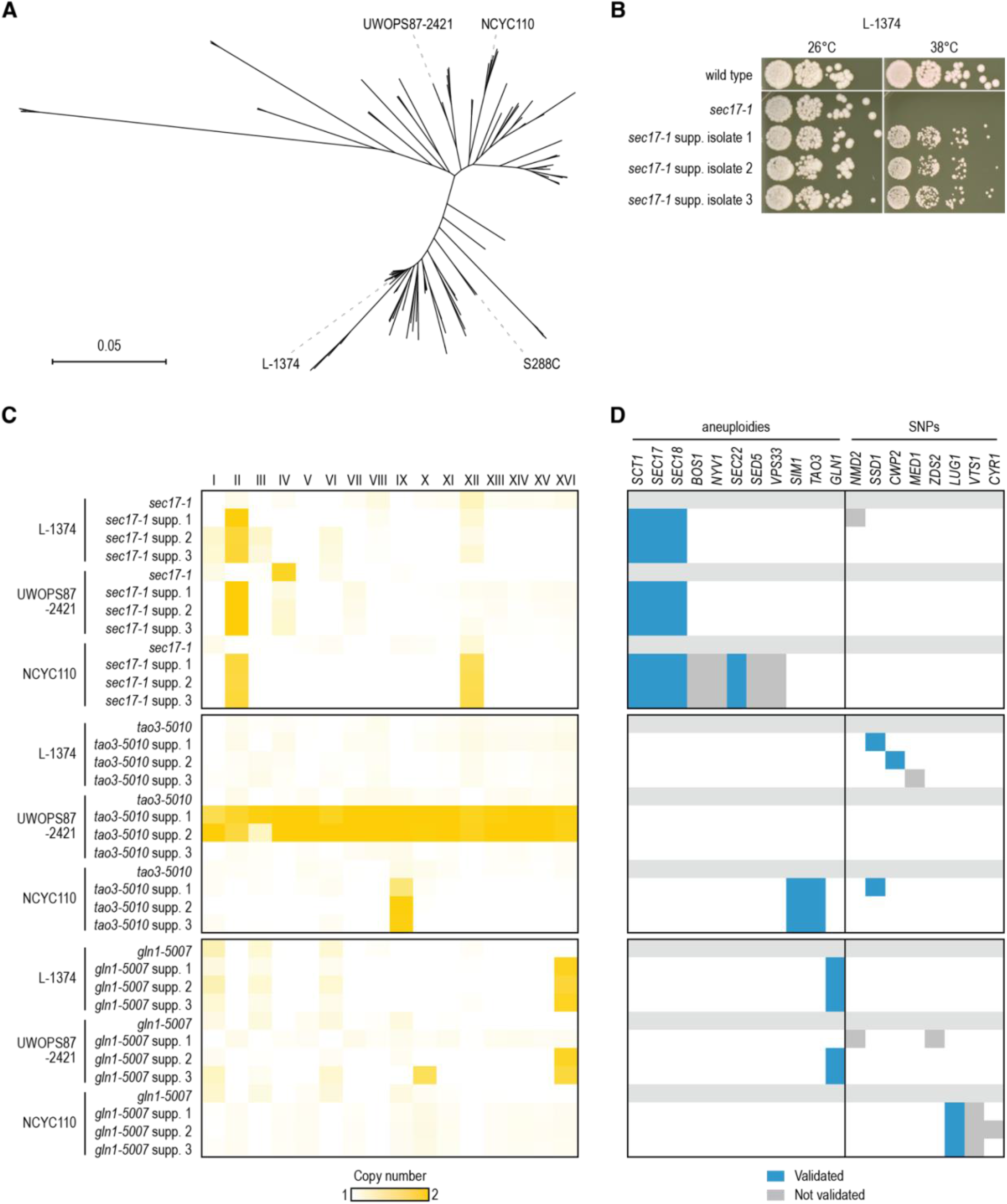
Systematic identification of genetic suppressors in diverse genetic backgrounds. (**A**) Phylogenetic tree of *Saccharomyces cerevisiae*, indicating the yeast isolates that were used in this study. Adapted from Batté et al., 2024. (**B**) Validating the suppression phenotype of isolated suppressor strains. Three TS alleles (*sec17-1, tao3-5010*, and *gln1-5007*) were introduced into three natural yeast isolates (L-1374, UWOPS87-2421, and NCYC110) and three independent, spontaneous suppressors of the TS phenotype were isolated in each background. An example of isolated suppressor strains of one TS allele (*sec17-1*) in one genetic background (L-1374) is shown here, all other combinations are shown in **Fig. S1**. Cultures of the isolated suppressor strains, as well as of the corresponding parental TS strain without a suppressor, were grown until saturation, and a series of ten-fold dilutions was spotted on YPD plates. Plates were incubated at the indicated temperatures for 2 days. The wild-type natural isolate (without TS allele) was included as a control. (**C**) All suppressor colonies, as well as the parental TS strains without suppressor, were sequenced whole genome. Shown is the average sequencing read depth for each of the 16 yeast chromosomes in each of the strains. Dark yellow indicates the presence of additional copies of the affected chromosome. (**D**) Validation of candidate suppressor genes. For candidate suppressor genes that were either located on one of the aneuploid chromosomes or that carried a nonsynonymous mutation, we tested the effect of deletion and/or overexpression of the genes on the temperature sensitivity of the query mutants. Blue squares indicate that the gene was found and validated as a suppressor of the TS allele in a given strain. Details on detected SNPs and aneuploidies can be found in **Data S1** and **S2**. Spot dilution assays of the suppressor validation experiments are shown in **Fig. 2, 3**, and **S2-S4** and results are summarized in **Data S3**.

We used the temperature sensitive phenotype of the constructed strains to isolate spontaneous suppressor mutants that could rescue the growth defect at high temperature. For each TS allele, we isolated three independent suppressor colonies per genetic background, for a total of 27 suppressor strains (**Fig. 1B, S1**). To identify the suppressor mutations, we sequenced the genomes of all 27 suppressor strains and the 9 corresponding parental strains. We identified 23 SNPs and 23 segmental or full aneuploidies that were present in a suppressor strain but not in the parental strain (**Fig. 1C**; **Data S1, S2**). Out of the 23 detected SNPs, 5 occurred in intergenic regions, 8 introduced premature stop codons or frameshifts that most likely led to loss of gene function, and 10 encoded missense variants. Most strains that carried nonsynonymous mutations did not carry aneuploidies, and vice versa. In total, 7 out of 27 suppressor strains were euploid and carried one or more nonsynonymous SNPs, 16 carried (partial) chromosomal duplications and no nonsynonymous SNPs, and 3 carried both a nonsynonymous SNP and an aneuploidy (**Data S3**). In the remaining suppressor strain, we could not identify any SNPs or other genomic alterations.

### Validating potential suppressor candidates

To determine which of the discovered genomic alterations were responsible for the suppression phenotype, we tested the effect of deletion and/or overexpression of the mutated genes on the temperature sensitivity of the query mutants (**Fig. 1D, S2-S4**; **Data S3**). In several cases, multiple suppressor strains carrying the same TS allele showed identical chromosome duplications (**Fig. 1C, Data S3**), suggesting that the suppression phenotype was caused by an increased copy number of one or more genes encoded on the affected chromosome. In 17 suppressor strains, the aneuploid chromosome carried the query TS allele itself, suggesting that increased dosage of the query allele caused the suppression. Indeed, transforming the parental TS strain (without the suppressor) with a plasmid carrying an extra copy of the TS allele improved fitness of all tested TS strains at elevated temperature (**Fig. 1D**; **Data S3**). However, in addition to the query allele itself, we suspected that in some cases other genes on the aneuploid chromosomes contributed to the suppression phenotype, as the fitness improvement caused by *sec17-1* and *tao3-5010* overexpression was modest in some backgrounds (**Fig. S2A, S3A**).

All *sec17-1* suppressor strains carried a duplication of chromosome II, which carries *sec17-1*. A previous study found that overexpression of either *SEC18* or *SCT1*, both located on chromosome II, could suppress *sec17-1* in S288C (Magtanong et al. 2011). We confirmed that overexpression of the S288C alleles of *SEC18* and *SCT1* could also suppress the *sec17-1* TS phenotype in the three natural genetic backgrounds (**Fig. 1D**; **Data S3**). Furthermore, the NCYC110 suppressor strains also carried a duplication of chromosome XII. Although there are no known dosage suppressors of *SEC17* located on this chromosome, it carries multiple genes with roles in vesicular transport. We tested five of these genes and found that only overexpression of *SEC22* could suppress the *sec17-1* TS phenotype (**Fig. 1D, S2C**; **Data S3**). Similarly, all NCYC110 *tao3-5010* suppressor strains carried an aneuploidy of chromosome IX, which carries *tao3-5010*. We validated that overexpression of *SIM1*, also located on chromosome IX and previously reported as a dosage suppressor of a *tao3* TS mutant in S288C (Du and Novick 2002), could suppress the *tao3-5010* TS allele in the NCYC110 background (**Fig. 1D**; **Data S3**).

To investigate a potential role for the identified nonsynonymous SNPs in the suppression phenotype, we introduced a plasmid carrying the wild-type alleles of the potential suppressor genes into the suppressor strains. If the suppressor mutation is recessive or semi-dominant, overexpression of the wild-type allele of the suppressor gene is expected to reverse the suppression and reduce the fitness of the suppressor strain. Using this strategy, we could not confirm a role for *NMD2, ZDS2, CYR1*, or *VTS1* in the suppression of *GLN1*, or for *MED1* in the suppression of *TAO3* (**Fig. 1D**; **Data S3**). However, expression of wild-type *SSD1* in L-1374 and NCYC110 *tao3-5010* suppressor strains carrying a missense variant in *SSD1* did revert the suppression phenotype, validating *SSD1* as the suppressor gene (**Fig. 1D**; **Data S3**). Furthermore, we deleted *SSD1* and *CWP2*, that carried potential loss-of-function variants in L-1374 and/or NCYC110 *tao3-5010* suppressor strains, in the parental strains that carry the TS allele but not the suppressor variant and confirmed that deletion of either of the genes could suppress *tao3* in these genetic backgrounds (**Fig. 1D**; **Data S3**). Similarly, we validated that deletion of *LUG1*, which carried loss-of-function variants in the NCYC110 *gln1-5007* suppressor strains, could suppress the temperature sensitivity of the parental NCYC110 *gln1-5007* strain (**Fig. 1D**; **Data S3**). Overall, we validated one or more suppressor genes in 22 out of 27 suppressor strains (**Fig. 1D**; **Data S3**).

### Suppression interactions are highly conserved across genetic backgrounds

In several cases, a particular suppressor gene was identified in only one genetic background. For example, *CWP2* was identified as a suppressor of *tao3-5010* only in the L-1374 background and loss-of-function mutations in *LUG1* were identified only in NCYC110 *gln1-5007* strains. However, these differences in observed suppressors across genetic backgrounds could be due to random chance or experimental factors. To directly investigate whether the identified suppressors were unique to a specific genetic context, we introduced deletion or overexpression alleles of the suppressors into all three natural strains, as well as in S288C, all carrying the query TS allele. For *GLN1*, we also tested for suppression by deletion of *PMR1*, a suppressor gene we had previously identified in the S288C background (our unpublished results) but not in any of the other backgrounds. Overexpression of *sec17-1, SEC18, SIM1*, or *gln1-5007* and deletion of *SSD1, CWP2*, or *LUG1* could suppress the corresponding query TS alleles in all four genetic backgrounds (**Table 1**; **Fig. S2-S4**). We did not succeed in deleting *PMR1* in the NCYC110 *gln1-5007* strain, but suppression was observed in the three remaining genetic backgrounds (**Table 1**; **Fig. S4E**). In contrast, overexpression of *tao3-5010* could suppress *tao3-5010* temperature sensitivity in the three natural backgrounds but not in S288C, possibly because of the high restrictive temperature of the allele in this genetic background (**Table 1**; **Fig. S3A**).

**Table 1.** Conservation of genetic suppression. For each of the detected suppressor alleles, its suppression phenotype was tested in all three natural yeast isolates, as well as in S288C. ✓ = suppression was observed in the indicated background; - = suppression was not observed in the indicated background; * = suppression was dependent on the expression level of the suppressor allele; n.d. = not determined; o.e. = overexpression. Spot dilution assays of the suppression assays are shown in **Fig. 2, 3**, and **S2-S4**.

| Query allele | Suppressor allele | S288C | L-1374 | UWOPS87-2421 | NCYC110 |
| --- | --- | --- | --- | --- | --- |
| *sec17-1* | *sec17-1* o.e. | ✓ | ✓ | ✓ | ✓ |
|  | *SEC18* o.e. | ✓ | ✓ | ✓ | ✓ |
|  | *SCT1* o.e. | ✓ | ✓ | ✓* | ✓* |
|  | *SEC22* o.e. | ✓ | ✓ | ✓* | ✓ |
| *tao3-5010* | *tao3-5010* o.e. | - | ✓ | ✓ | ✓ |
|  | *ssd1*Δ | ✓ | ✓ | ✓ | ✓ |
|  | *cwp2*Δ | ✓ | ✓ | ✓ | ✓ |
|  | *SIM1* o.e. | ✓ | ✓ | ✓ | ✓ |
| *gln1-5007* | *gln1-5007* o.e. | ✓ | ✓ | ✓ | ✓ |
|  | *lug1*Δ | ✓ | ✓ | ✓ | ✓ |
|  | *pmr1*Δ | ✓ | ✓ | ✓ | n.d. |

For *SCT1* and *SEC22*, that could both suppress *sec17-1*, suppression was dependent on the expression level of the suppressor gene. A *CEN*-plasmid (low-copy) containing *SCT1* could suppress *sec17-1* in S288C and L-1374, but a *2µ*-plasmid (high-copy) was needed to see suppression in the NCYC110 or UWOPS87-2421 backgrounds (**Fig. 2A**). Because we were using S288C alleles in the overexpression experiments, we tested whether overexpression of the *SCT1* allele from NCYC110 or UWOPS87-2421 could suppress *sec17-1* in these backgrounds when expressed from a low-copy plasmid. However, also the NCYC110 and UWOPS87-2421 *SCT1* alleles were not able to suppress *sec17-1* in these backgrounds when expressed from a *CEN*-plasmid (**Fig. 2B**). Similarly, overexpression of *SEC22* from a *CEN*-plasmid could rescue *sec17-1* strains in most genetic backgrounds, but not in UWOPS87-2421 (**Fig. 3A**). In this case, further increasing the level of overexpression of the *SEC22* S288C did not rescue *sec17-1* in the UWOPS87-2421 background (**Fig. 3A**), but overexpression of the *SEC22* UWOPS87-2421 allele showed weak but reproducible suppression (**Fig. 3B**). Although the sequence of the *SEC22* ORF is identical in UWOPS87-2421 and S288C, the UWOPS87-2421 allele contains a C-to-T variant in the 5’ UTR, 69 nucleotides upstream of the start codon. We investigated the effect of this UWOPS87-2421-specific variant on *SEC22* mRNA levels using RNA sequencing and found an ∼25% increase in *SEC22* expression in the wild-type UWOPS87-2421 strain compared to S288C (**Fig. 3C**; **Data S4**). Possibly, suppression of *sec17-1* by *SEC22* in the UWOPS87-2421 background is sensitive to small changes in *SEC22* expression. Expressing the S288C *SEC22* allele from a *CEN-* or *2µ*-plasmid may lead to too low or too high expression, respectively, while expression of the UWOPS87-2421 *SEC22* allele from a *CEN* plasmid may achieve a level of expression that is just right for suppression to occur.

**Figure 2.**
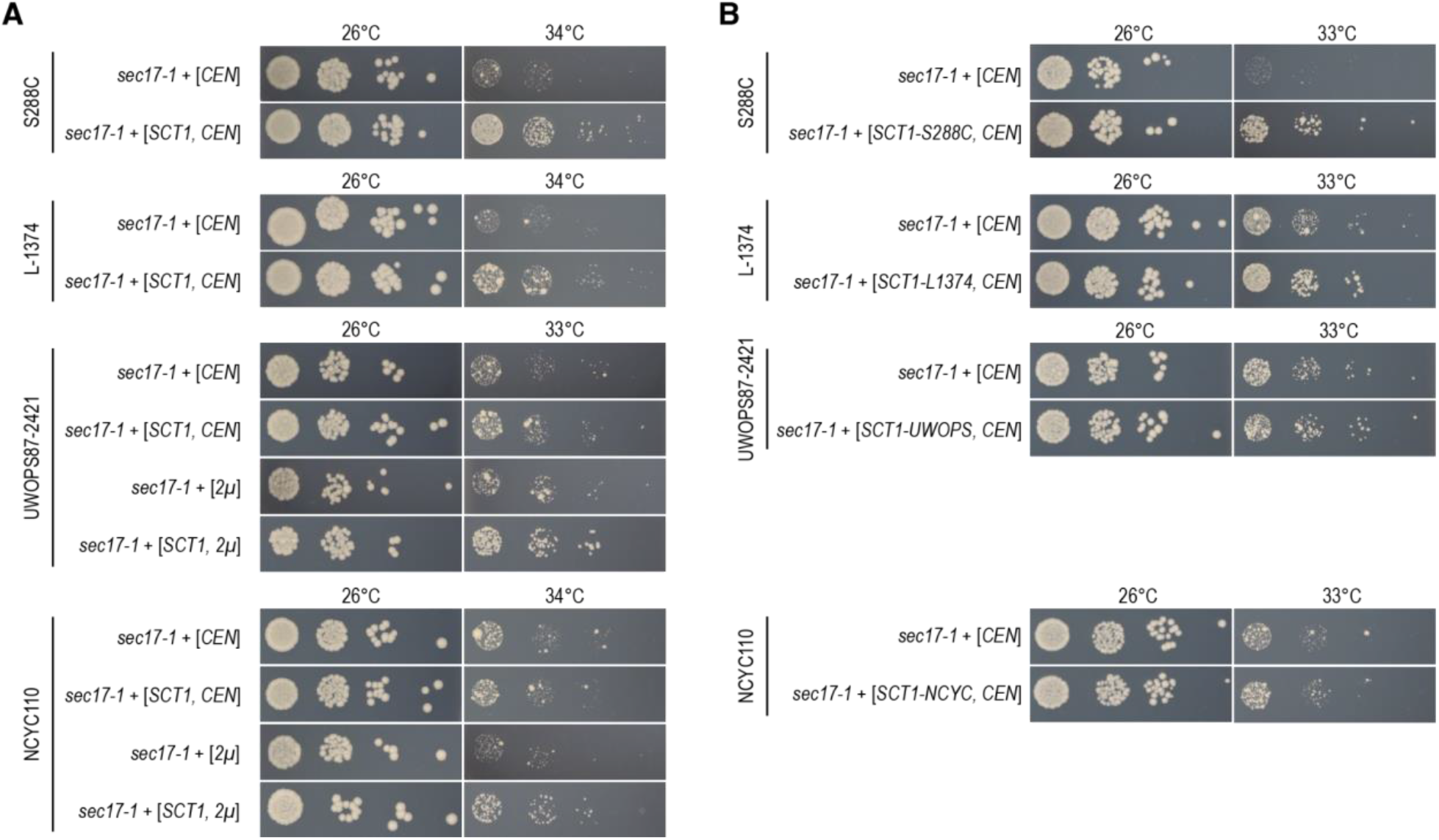
Different levels of *SCT1* expression are needed for suppression of *SEC17* across genetic backgrounds. (**A**) S288C, L-1374, UWOPS87-2421, and NCYC110 strains carrying the *sec17-1* TS allele were transformed with a low-copy (*CEN*) or a high-copy (*2µ*) plasmid expressing *SCT1* or the corresponding empty vector. Cultures of two to three independent transformants were grown until saturation, and a series of ten-fold dilutions was spotted on SD-Ura (low-copy) or SD-Leu (high-copy) plates. Plates were incubated at the indicated temperatures for 3 days. Pictures of one representative transformant are shown for each genotype. Rare, larger colonies that appear at higher temperatures are spontaneous suppressor mutants that sometimes occur during the experiments. (**B**) As in (A), but using the *SCT1* alleles from the L-1374, UWOPS87-2421, and NCYC110 backgrounds, rather than the S288C allele. UWOPS = UWOPS87-2421; NCYC = NCYC110.

**Figure 3.**
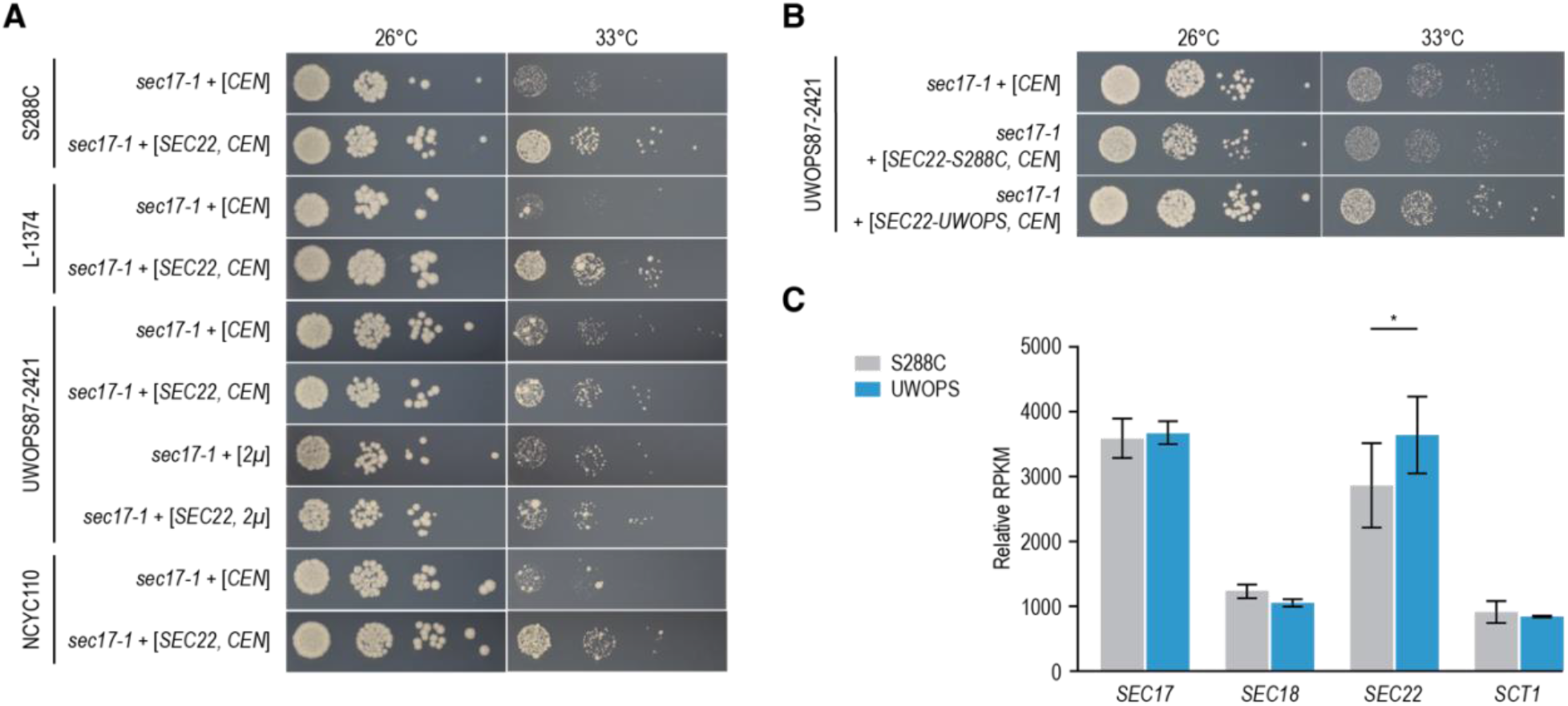
Suppression of *SEC17* by *SEC22* is dependent on the allele sequence. (**A**) S288C, L-1374, UWOPS87-2421, and NCYC110 strains carrying the *sec17-1* TS allele were transformed with a low-copy (*CEN*) or a high-copy (*2µ*) plasmid expressing *SEC22* or the corresponding empty vector. Cultures of two to three independent transformants were grown until saturation, and a series of ten-fold dilutions was spotted on SD-Ura (low-copy) or SD-Leu (high-copy) plates. Plates were incubated at the indicated temperatures for 3 days. Pictures of one representative transformant are shown for each genotype. (**B**) A UWOPS87-2421 strain carrying the *sec17-1* TS allele was transformed with a low-copy vector carrying either the S288C or the UWOPS87-2421 version of *SEC22*, or the corresponding empty vector. Spot dilutions were performed as in (A). (**C**) Expression levels of the indicated genes in wild-type S288C or UWOPS87-2421 strains were determined by RNA sequencing. Plotted are RPKM (reads per kilobase per million mapped reads) values, normalized to the total number of reads in a sample and averaged over three technical replicates. Error bars indicate the standard deviation. * p<0.05, two-sided Student’s t-test. UWOPS = UWOPS87-2421.

Thus, 10 out of 11 suppression mechanisms tested in this study (8 out of 8 when excluding suppression by overexpression of the query allele) were conserved in all tested genetic backgrounds, with only the required expression level of the suppressor gene changing with the genetic context. Despite the high conservation of suppressor genes, the relative strength of the suppressors varied between genetic backgrounds. For example, overexpression of *SIM1* could strongly suppress *tao3-5010* in the NCYC110 and L-1374 backgrounds, but only weakly improved fitness in the S288C and UWOP87-2421 backgrounds (**Fig. S3B**). Such differences in the intensity of suppression across genetic backgrounds were common and observed for nearly all suppressor genes. A notable exception is *ssd1*Δ, which could strongly suppress *tao3-5010* in all backgrounds (**Fig. S3E**).

### Multiple genes can contribute to the suppression phenotype

In a few instances, we had identified multiple genes on aneuploid chromosomes that could each independently suppress the TS phenotype (**Data S3**). We hypothesized that in these cases the suppressors could have an additive effect, and that the combined overexpression of multiple suppressor genes may further improve the fitness of the TS mutant at higher temperature. To test this, we combined overexpression of *SEC18, SEC22*, and *SCT1* in *sec17-1* strains. *SEC18* and *SCT1* are both located on chromosome II, which was duplicated in all *sec17-1* suppressor strains, and *SEC22* is located on chromosome XII, which was duplicated together with chromosome II in the NCYC110 *sec17-1* suppressor strains. We constructed a collection of *CEN*-plasmids, each expressing a natural allele of one of the three genes and a different selectable marker and verified that each gene individually could suppress *sec17-1* in the same backgrounds as described above (**Fig. S5A**). We then overexpressed all possible combinations of the three suppressors in their respective genetic background in presence of the *sec17-1* TS allele (**Fig. 4, S5B**). To be able to detect small differences in fitness, we quantified the size of on average ∼200 colonies per strain (**Fig. 4**).

**Figure 4.**
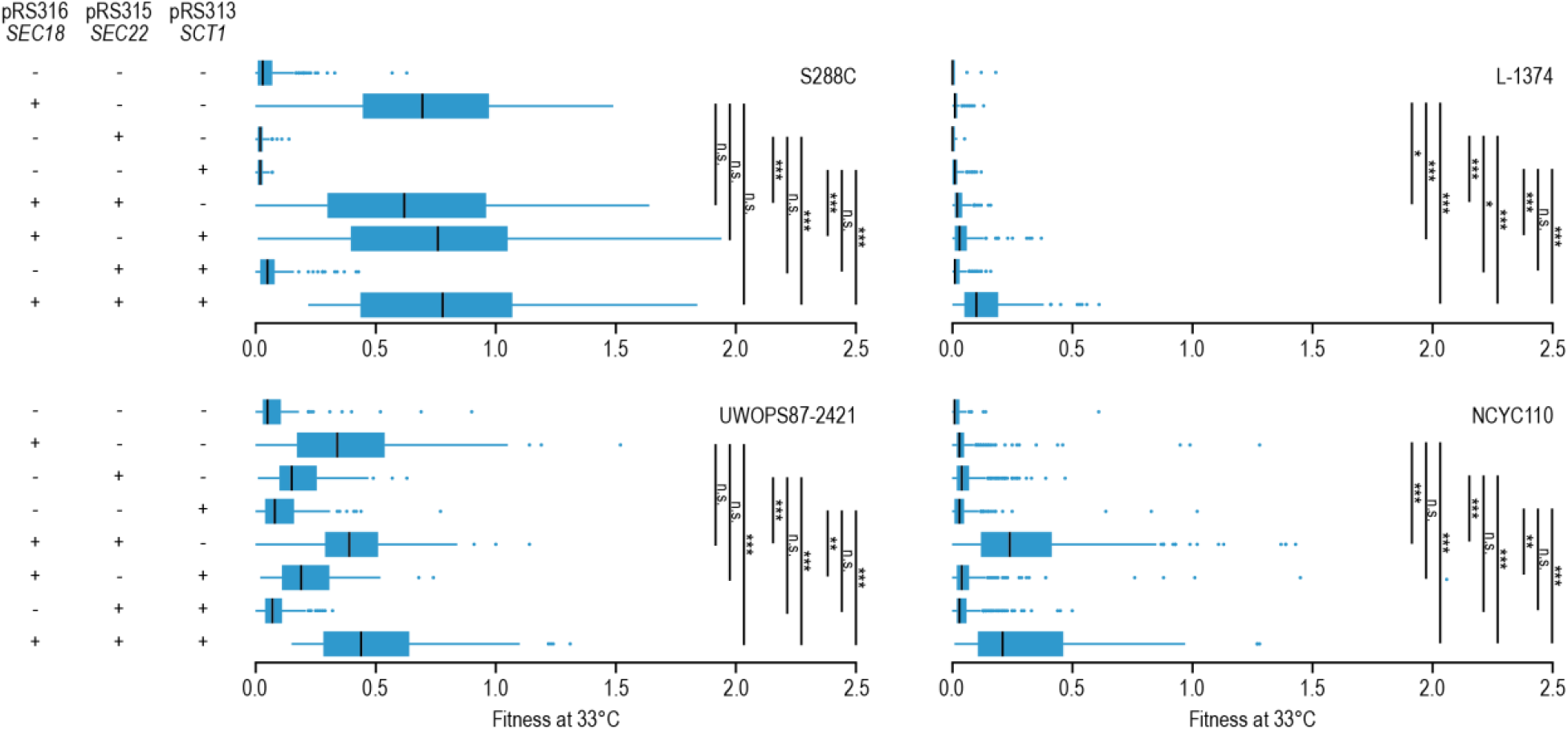
Multiple genes can contribute to the suppression phenotype. Relative fitness of *sec17-1* strains overexpressing *SEC18, SEC22*, and/or *SCT1* in the S288C, L-1374, UWOPS87-2421, and NCYC110 genetic backgrounds. In each case, the *SEC18, SEC22*, and *SCT1* alleles matched the genetic background in which they were transformed, such that S288C was transformed with S288C alleles and L-1374 with L-1374 alleles, etc. Strains were spotted on SD-Leu-Ura-His and colony size was determined after two days of growth at 33°C and normalized to the colony size of a wild-type strain with the same genetic background. Boxplots show the normalized colony size (fitness) of, on average, ∼200 colonies per strain. Tukey’s test was used to determine whether strains overexpressing two or three genes had a significantly higher fitness than strains overexpressing one of the genes. + = strains were transformed with the indicated plasmid; - = strains were transformed with the corresponding empty vector. * p<0.05; ** p<0.005; *** p<0.0005.

In S288C and UWOPS87-2421, the suppression was mainly driven by the strongest suppressor of *sec17-1, SEC18*, and little further increase in suppression was observed when *SCT1* and/or *SEC22* were overexpressed simultaneously with *SEC18* (**Fig. 4**). In contrast, in L-1374, overexpression of all three genes simultaneously resulted in stronger suppression than overexpression of each of the genes alone (**Fig. 4**). Combining *SEC18* (chromosome II) and *SEC22* (chromosome XII) overexpression strongly improved fitness compared to overexpression of *SEC18* or *SEC22* alone in the NCYC110 *sec17-1* strain, which is also the genetic background in which combined duplication of chromosome II and XII was observed. These results show that multiple genes can contribute to the suppression phenotype caused by aneuploidies, and that the relative contribution of the individual genes to the overall suppression varies between genetic backgrounds.

## DISCUSSION

In this study, we investigated the conservation of genetic suppression interactions across natural yeast isolates using three mutant alleles of functionally diverse query genes. Ten out of eleven mechanisms of suppression that spontaneously occurred in the natural yeast strains could be reproduced in all four tested genetic backgrounds, including the laboratory strain S288C. Despite the high conservation of suppression interactions, the extent of suppression was often variable between backgrounds, and sometimes depended on the expression level (**Fig. 2**) or allele sequence (**Fig. 3**) of the suppressor gene. Similarly, a previous study compared five suppressor genes of a *las17* TS allele in the yeast strains S288C and RM11-1a and found that the strength of the suppression phenotype varied between the two strains and was in some cases influenced by the particular suppressor mutation (Filteau et al. 2015). Some of the differences in strength and required sequence of the suppressor genes could be due to differences in the restrictive temperature of the TS allele between genetic backgrounds. A large difference in restrictive temperature may also explain why the *tao3-5010* mutant could not be rescued by overexpression of the TS allele in S288C, in which the mutant had a restrictive temperature of 38°C, in contrast to the other genetic backgrounds where the restrictive temperature was ∼8-10 degrees lower (**Fig. S3A**). Possibly, the remaining functionality of the *tao3-5010* allele at 38°C is insufficient to support proliferation. Alternatively, these differences may result from strain-specific variants in additional genes.

Most of the isolated suppressor strains in the natural backgrounds carried aneuploidies (19 out of 27, 70%), and in 17 out of 19 cases we validated that one or more genes on the aneuploid chromosome were responsible for the suppression phenotype (**Data S3**). This frequency of aneuploidies is significantly higher than what we generally observe for suppressors of TS alleles in S288C (∼16% carry aneuploidies, our unpublished results). We suspect that this difference in aneuploidy occurrence is due to natural yeast strains being relatively tolerant to aneuploidies when compared to S288C (Hose et al. 2015). Aneuploidies are associated with a growth defect in laboratory yeast strains, independently of which chromosome is duplicated (Torres et al. 2007, Beach et al. 2017). In contrast, the sequencing of more than a thousand yeast isolates showed that ∼20% of natural *S. cerevisiae* strains carry at least one aneuploidy (Peter et al. 2018). Because aneuploidies commonly occur during cell division (Gilchrist and Stelkens 2019), the enhanced tolerance for aneuploidies may increase the frequency at which suppression occurs, which could be an advantage in highly selective natural environments. Furthermore, we showed that multiple genes on an aneuploid chromosome can contribute to the suppression phenotype (**Fig. 4**), further increasing the benefit associated with an aneuploidy.

Out of the eight extragenic suppressors that we identified in this study, four had not been described previously. For example, we found that overexpression of SNARE protein Sec22 could suppress mutants of the SNARE chaperone Sec17 (**Fig. 3**), likely by (partially) restoring vesicle fusion (Liu and Barlowe 2002, Song et al. 2021). Furthermore, we discovered loss-of-function mutations in *CWP2*, encoding a major cell wall mannoprotein, as suppressors of the RAM signaling network member Tao3 (**Fig. S3E**). Cells with an inactive RAM network display a separation defect of mother and daughter cell walls due to the inability to activate Ace2, which is needed for the expression of cell separation genes (Nelson et al. 2003). Possibly, the changes in cell wall composition induced by loss of *CWP2* (Van der Vaart et al. 1995, Li et al. 2020) can promote the separation of mother and daughter cells in the absence of a functioning RAM network. We also found that loss-of-function mutations in *PMR1* could suppress a TS mutant of the glutamine synthetase Gln1 in all tested genetic backgrounds (**Fig. S4E**). Pmr1 shuttles calcium and manganese (Mn^2+^) ions into the Golgi lumen, and loss of Pmr1 leads to increased intracellular levels of Mn^2+^ (Lapinskas et al. 1995, Durr et al. 1998). Glutamine synthetases are activated by Mn^2+^ ions (Monder 1965, Tholey et al. 1987), suggesting that loss of *PMR1* may suppress the *GLN1* mutant by boosting its activity. Finally, we uncovered loss-of-function mutations in the poorly characterized *LUG1* gene as suppressors of *GLN1* (**Fig. S4D**). Mutations in *GLN1* were previously described to suppress *lug1*Δ mutants, indicating that this suppression interaction is reciprocal (Edskes et al. 2018). Overall, this high frequency of newly identified suppressor genes indicates that despite several large-scale suppressor mapping efforts (Magtanong et al. 2011, Patra et al. 2016, Van Leeuwen et al. 2016, Van Leeuwen et al. 2020), the yeast suppression interaction space remains largely unexplored.

In conclusion, although different genetic backgrounds have the potential to reveal novel suppression interactions and thus uncover previously unidentified functional connections between genes (Filteau et al. 2015), our results suggest that genetic suppression interactions are largely robust to changes in genetic context. While the extent of conservation of suppression interactions across human genomes remains to be determined, our finding is potentially reassuring for the development of new therapeutic strategies that target suppressor genes (Esrick et al. 2021, Frangoul et al. 2021, Ünlü et al. 2023).

## METHODS

### Yeast strains, plasmids and growth

Yeast strains were grown using standard rich (YPD) or minimal (SD) media. For overexpression assays using S288C alleles, plasmids from either the MoBY-ORF 1.0 (native promoter, *CEN/ARS, URA3, kanMX4*) (Ho et al. 2009) or the MoBY-ORF 2.0 (native promoter, 2μ, *LEU2, kanMX4*) (Magtanong et al. 2011) collection were used. All yeast strains and plasmids used in this study are listed in **Data S5**.

### Introducing TS alleles into multiple genetic backgrounds

The three natural yeast isolates, L-1374 (LY00010), UWOPS87-2421 (LY00011), and NCYC110 (LY00015), were previously (partially) deleted for *HO, URA3, HIS3*, and *LEU2*, resulting in the genotype MATa *ho*Δ*::hphMX6 ura3*Δ*::kanMX his3*Δ*1 leu2*Δ*0* (Cubillos et al. 2009, Parts et al. 2021). The three genetically diverse strains were each crossed with three different S288C strains carrying a TS allele, TSQ48 (*sec17-1*), TSQ2433 (*gln1-5007*), and TSQ2031 (*tao3-5010*), with genotype MATα *xxx-ts::natMX4 can1*Δ*::STE2pr-Sp_his5 lyp1*Δ*0 his3*Δ*1 leu2*Δ*0 ura3*Δ*0 met15*Δ*0*. The resulting diploids were driven through meiosis and haploid MATα segregants carrying the TS allele were isolated and crossed again to their respective natural parental strain. This process was repeated four more times for a total of six crosses per strain background and TS allele, resulting in progeny with a genome that is 98% identical to the natural parental strain. Two independent spores carrying the TS allele were isolated from the final crosses and frozen at -80°C.

The S288C and L-1374 *sec17-1* strains carried the *lyp1*Δ*::STE3pr-LEU2* cassette, complicating some of the suppressor validation experiments that used *LEU2*-plasmids. To remove the *lyp1*Δ*::STE3pr-LEU2* cassette, we first cloned *LEU2*-targeting guide RNA sequences into the pML104 vector, which carries Cas9 and a *URA3* selection marker (**Data S5**) (Laughery et al. 2015). Next, we PCR-amplified *LYP1*, including promoter and terminator sequences, from a wild-type strain and co-transformed the *lyp1*Δ*::STE3pr-LEU2* strains with the *LYP1* PCR-product and the pML104-LEU2-2 plasmid. Transformants were selected on SD-Ura and subsequently streaked on SD-Leu and SD-Lys+LYP (thialysine) to confirm loss of *LEU2* and proper integration of *LYP1*. The final genotypes and strain IDs of the resulting strains are listed in **Data S5**.

### Isolating spontaneous suppressor mutations

For each TS allele in each genetic background, ∼25 million cells were spread onto three YPD+NAT agar plates and incubated for three days at the restrictive temperature of the strain. Most cells will not be able to grow at the restrictive temperature, except for those that have acquired a spontaneous suppressor mutation, that will grow up to form a colony. When colonies were observed, one colony per plate was isolated and its growth at the restrictive temperature was compared to the parental TS strain to confirm the suppression phenotype. In total, three independent suppressors per query allele and per genetic background were isolated.

### Sequencing, read mapping, and SNP calling

All suppressor strains as well as the corresponding parental TS strains were sequenced on the DNBseq platform using paired-end 100-bp reads, with an average read depth of ∼100x. Reads were aligned to the S288C reference genome version R64.2.1 using BWA v0.7.17 (Li and Durbin 2009). Pileups were processed and variants were called using SAMtools/BCFtools v1.11 (Li et al. 2009). Variants that had a Phred quality score <200, that were present in one of the parental strains, or that were found in more than three of the suppressor strains were removed from consideration. The consequence of detected variants was determined using Ensembl’s VEP (McLaren et al. 2016). All whole-genome sequencing data are publicly available at NCBI’s Sequence Read Archive (http://www.ncbi.nlm.nih.gov/sra) under accession number PRJNA1100912. Variants are listed in **Data S1**.

### Aneuploidy and ploidy assessment

Qualimap v2.3 (Okonechnikov et al. 2016) was used to detect (partial) aneuploidies based on variation in sequencing read depth across windows of 30,000 base pairs in the nuclear genome (**Data S2**). We note that the smaller chromosomes I, III, and VI showed a higher variation in read count between samples than other chromosomes, likely due to variation in the capture of these small chromosomes during genomic DNA isolation. Because the relative increase in coverage caused by an aneuploidy depends on the overall ploidy, we analyzed all suppressor strains by flow cytometry to determine ploidy. Briefly, cells were grown until log-phase (OD_600_≈0.5) and fixed with 70% ethanol. Fixed cells were washed with water and subsequently treated with RNase A (200 µg/ml, 2 h, 37°C) and proteinase K (2 mg/ml, 40 min, 50°C). Treated cells were washed with 200 mM Tris–HCl, 200 mM NaCl, 78 mM MgCl2 (pH 7.5) and stained with 2× SYBR Green (Life Technologies) in 50 mM Tris–HCl (pH 7.5). Aggregates of cells were dispersed via sonication and cells were analyzed by flow cytometry using a SONY SH800 FACS machine. DNA content was compared to known haploid and diploid controls. Normalized average read depth per genomic region was corrected based on the observed DNA content, such that the average normalized read depth of a genomic region in a diploid strain was twice that of a haploid strain. Detected aneuploidies are summarized in **Data S3**.

### Predicting and validating suppressor genes

For suppressor strains that carried an aneuploidy, we predicted potential causal suppressor genes based on the functional relationships between the query gene and the genes located on the aneuploid chromosome. We used BioGRID 4.4 (Oughtred et al. 2021) to identify genes that are known to interact with the query gene (either genetically or physically) and the *Saccharomyces* Genome Database (Wong et al. 2023) to identify genes that function in similar or related biological processes as the query. Identified candidate suppressors were validated by transforming plasmids expressing the candidate gene into the parental TS strain, without the suppressor, using standard transformation protocols (Gietz and Schiestl 2007). Overnight cultures of three independent transformants were diluted to an OD_600_ of 0.1, serially diluted 1:10 with sterile water, and spotted onto agar plates. Plates were incubated at a range of temperatures between 26°C and 38°C. After 2-3 days of incubation, pictures were taken and the relative fitness of the transformants was compared to empty vector controls.

To test whether detected nonsynonymous SNPs contributed to the suppression phenotype, we introduced a plasmid carrying the wild-type allele of the potential suppressor gene into the suppressor strain. If the suppressor mutation is recessive or semi-dominant, overexpression of the wild-type allele of the suppressor gene is expected to reverse the suppression and reduce the fitness of the suppressor strain. Transformations and spot dilutions assays were performed as described above for the aneuploidy suppressors. Validated suppressor genes are listed in **Data S3**.

### Suppression by overexpression of natural alleles

To test for suppression by overexpression of the natural alleles of suppressor candidates, we constructed plasmids carrying these alleles. We PCR-amplified the suppressor candidates including ∼1000 bp upstream of the start codon and ∼500 bp downstream of the stop codon from the various natural yeast strains, thereby including regions of homology to plasmid pRS313, pRS315, or pRS316 (Sikorski and Hieter 1989) (**Data S5**). The PCR product was co-transformed with the corresponding linearized vector into LY00004 (BY4742; **Data S5**). The assembled plasmid was isolated from the yeast strain and correct insertion of the PCR product was verified using whole plasmid sequencing. Plasmids were transformed into parental TS strains and tested for suppression as described above (“*Predicting and validating suppressor genes”*).

### Validation of SSD1, CWP2, PMR1, and LUG1

To test whether deletion of *SSD1* or *CWP2* could suppress *tao3* TS alleles in the S288C genetic background, *ssd1*Δ (DMA1035; **Data S5**) and *cwp2*Δ (DMA2828; **Data S5**) strains were crossed to *tao3-5005* (TSQ2026; **Data S5**) and *tao3-5014* (TSQ2035; **Data S5**). Diploids were selected on YPD+NAT+G418, driven through meiosis, and haploid double mutant progeny were isolated using tetrad dissection. In total, three independent double mutant spores were isolated per cross.

To test whether deletion of *SSD1* or *CWP2* could suppress *tao3* TS alleles in the natural genetic backgrounds, we PCR-amplified *CaURA3MX4* from plasmid pFA6:CaURA3MX4 (Goldstein et al. 1999) (**Data S5**), thereby introducing regions of homology to the genomic DNA directly upstream and downstream of both suppressor genes. The PCR products were transformed into the natural *tao3-5010* strains and deletion of *CWP2* and *SSD1* was verified by PCR. A similar strategy was used to delete *PMR1* and *LUG1* in the *gln1-5007* strains, with the exception that we cloned a guide RNA targeting *PMR1* or *LUG1* into the Cas9-expressing vector pML107 (Laughery et al. 2015). We then co-transformed the cloned plasmids with the *CaURA3MX4* cassettes to increase the efficiency of gene deletion. All strains were tested for suppression as described above (“*Predicting and validating suppressor genes”*).

### RNA sequencing

Overnight cultures of S288C and UWOPS87-2421 were diluted in 10 mL YPD to an OD_600_ of 0.1 and grown for 3-4 hours at 26°C until an OD_600_ of ∼0.7-1.0. Cells were collected, washed with water, snap frozen in liquid nitrogen, and stored at -80°C until RNA extraction. Total RNA was extracted by first lysing the yeasts with glass beads in trizole, separating the protein-DNA-RNA phases with chloroform, and precipitating the RNA with isopropanol and glycogen. The resulting RNA was washed with 70% ethanol, dissolved in water, treated with DNAse, and further cleaned using the Macherey-Nagel NucleoSpin RNA kit. RNA quality was assessed using a Fragment Analyzer and mRNA was enriched via polyA-selection with the Illumina Stranded mRNA Prep kit and sequenced on the Element Biosciences AVITI system using 150 base pair, single-end reads with ∼20 million reads per sample. Adapters were trimmed from the reads with Cutadapt v2.5 (Martin 2011) and reads with low complexity sequences were removed with Reaper v15-065 (Davis et al. 2013). Reads corresponding to ribosomal RNAs were removed with FastQ Screen v0.11.1 (Wingett and Andrews 2018). Remaining reads were aligned with STAR v2.5.3a (Dobin et al. 2013) against reference genome R64.2.1. The number of read counts per gene locus was summarized with HTSeq-count v0.9.1 (Anders et al. 2015) and normalized to gene length and the total number of reads per sample. Normalized read counts are listed in **Data S4**.

### Quantifying strain fitness

To determine the fitness of strains overexpressing combinations of SEC18, SEC22, and/or SCT1, saturated cultures of three independent transformants per genotype were diluted 1,000 to 100,000 fold and spotted onto SD-His-Leu-Ura plates. Plates were incubated at 26°C or 33°C and imaged after two days. All images were edited in an identical way to achieve maximal sharpness and to increase contrast by 10% and highlights by 5%. Images were then cropped, and colony size was determined using CellProfiler version 4.2.8 (Carpenter et al. 2006). Colony sizes were normalized to the median colony size of the wild-type strain of the same genetic background.

## DATA AVAILABILITY

All whole-genome sequencing data are publicly available at NCBI’s Sequence Read Archive (http://www.ncbi.nlm.nih.gov/sra) under accession number PRJNA1100912. All other data are available in the supplementary data files.

## ACKNOWLEDGEMENTS

We thank Sabine van Schie for critical reading of the manuscript. This work was supported by an Eccellenza grant from the Swiss National Science Foundation (PCEGP3_181242 to J.v.L).

