## Supplementary Figures for "Genetic suppression interactions are highly conserved across genetic backgrounds"

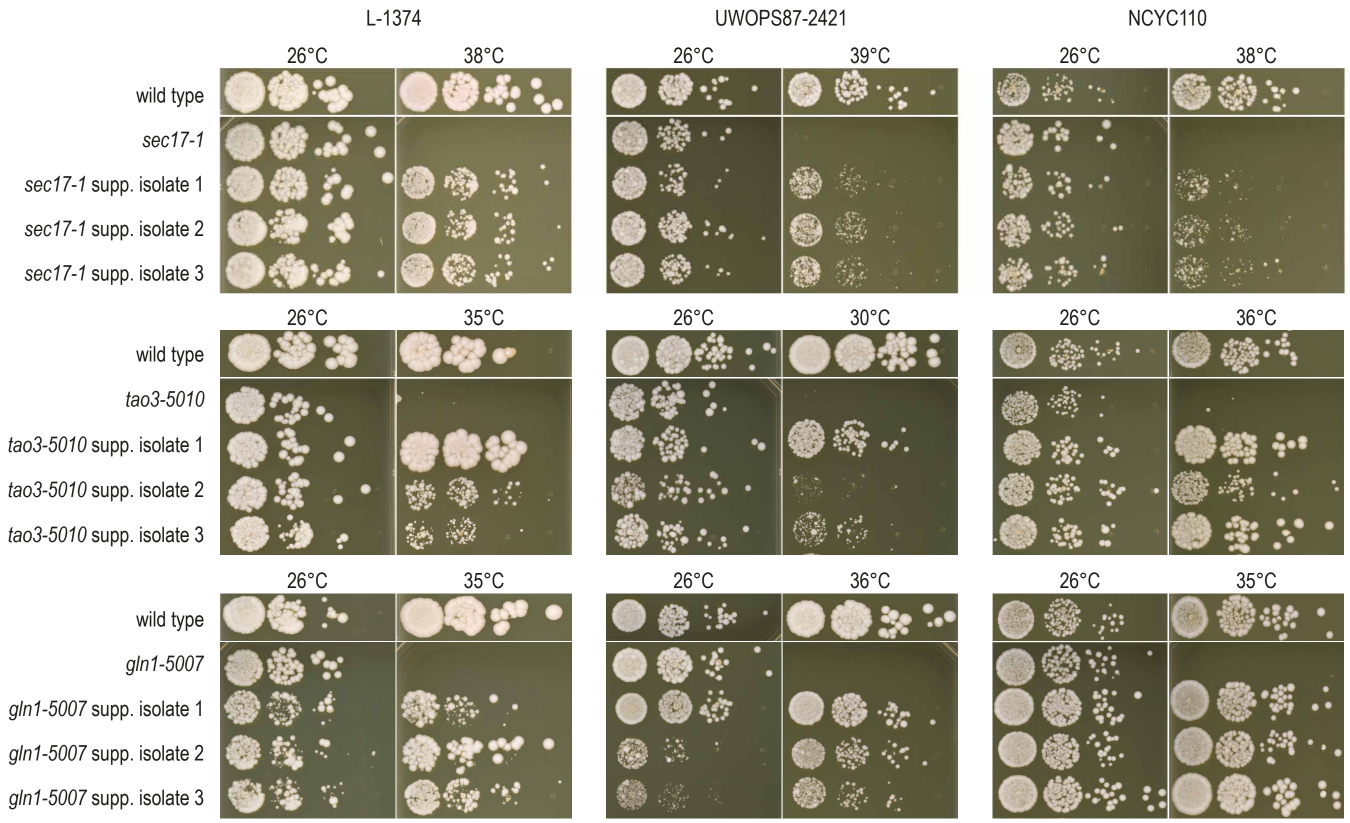


**Fig. S1. Validating the suppression phenotype of isolated suppressor strains.** Three TS alleles (*sec17-1*, *tao3-5010*, and *gln1-5007*) were introduced into three natural yeast isolates (L-1374, UWOPS87-2421, and NCYC110) and spontaneous suppressors of the TS phenotype were isolated. Cultures of the isolated suppressor strains, as well as of the corresponding parental TS strains without a suppressor, were grown until saturation, and a series of ten-fold dilutions was spotted on YPD plates. Plates were incubated at the indicated temperatures for 2 days. The wild-type natural isolates (without TS allele) were included on each plate as a control.

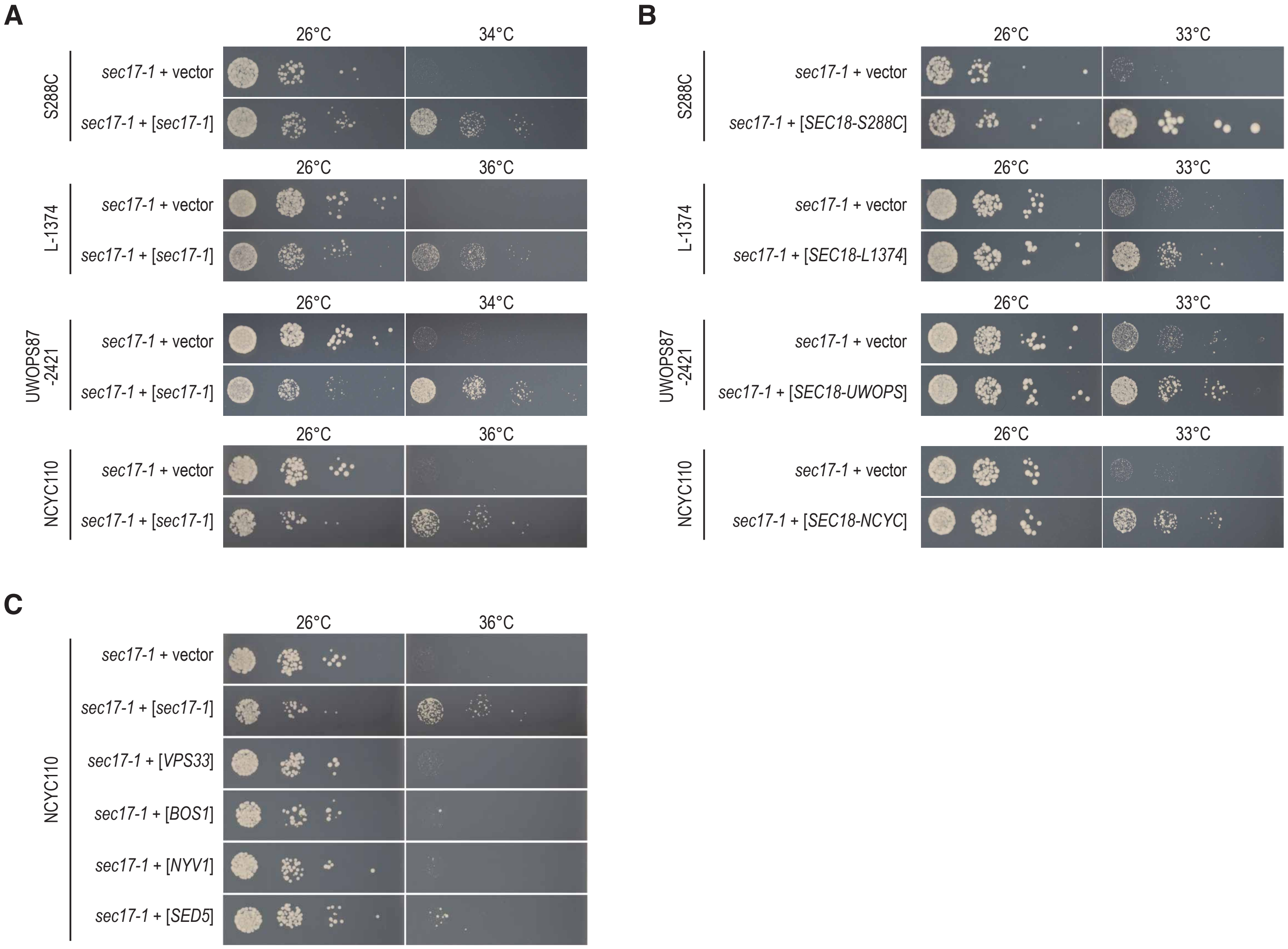


**Fig. S2. Validation of *SEC17* suppressors.** (**A**-**C**) Suppression of *sec17-1* TS strains by overexpression of genes located on aneuploid chromosomes: *sec17-1* (A) or *SEC18* (B), both located on chromosome II, or genes located on chromosome XII (C). Cultures of three independent transformants of the indicated strains were grown until saturation, and a series of ten-fold dilutions was spotted on SD-Ura plates. Plates were incubated at the indicated temperatures for 2 days. Pictures of one representative transformant are shown for each genotype. UWOPS = UWOPS87-2421; NCYC = NCYC110.

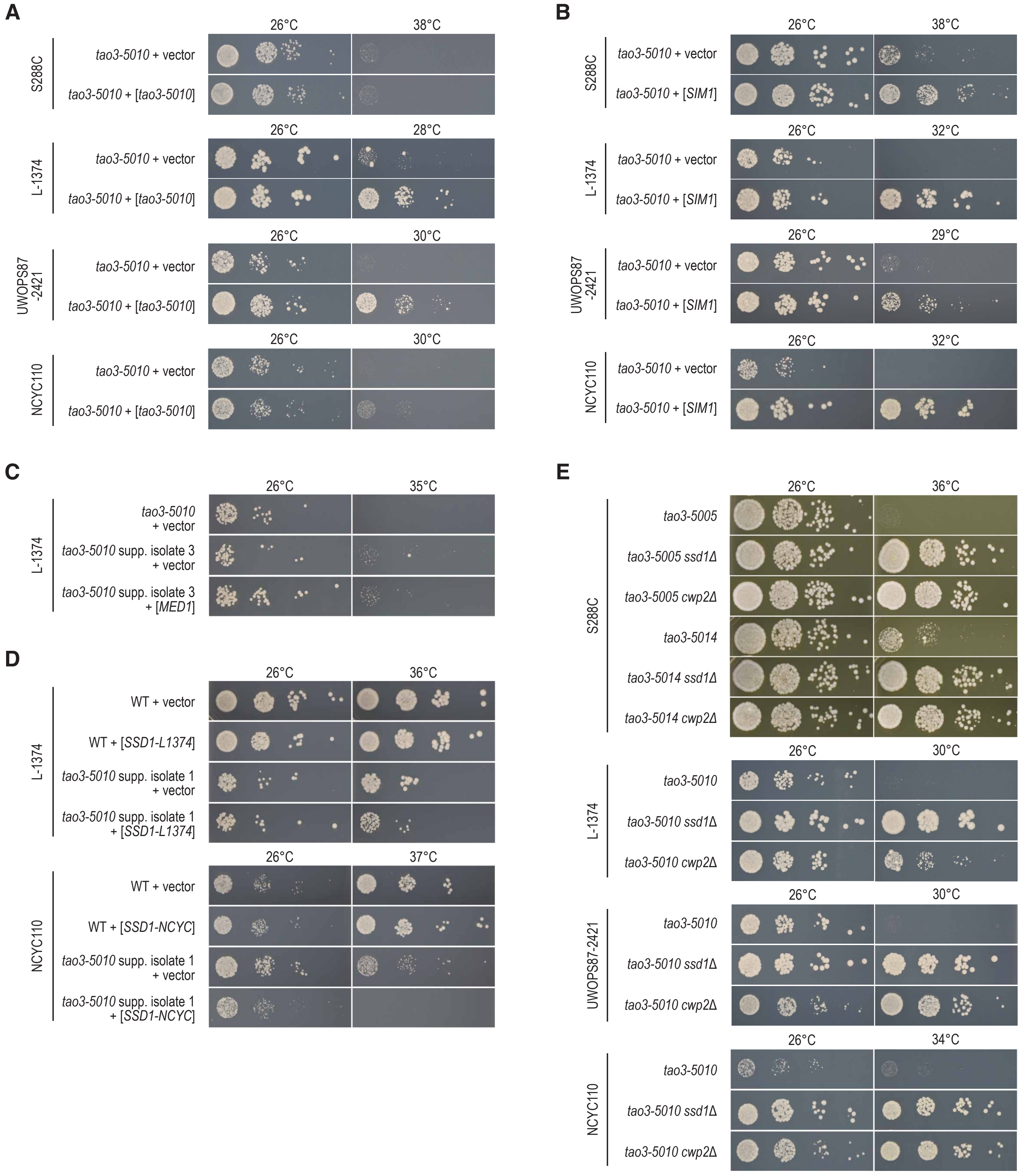


**Fig. S3. Validation of *TAO3* suppressors.** (**A-E**) Spot dilution assays of *tao3-5010* strains overexpressing *tao3-5010* (A), *SIM1* (B), *MED1* (C), or *SSD1* (D), or deleted for *SSD1* or *CWP2* (E). Cultures of 2-3 independent isolates of the indicated strains were grown until saturation, and a series of ten-fold dilutions was spotted on SD-Ura+NAT (A: S288C), SD-Leu (B), SD-Ura (A and E: L-1374, UWOPS87-2421, NCYC110; C; D), or YPD+NAT (E: S288C). Plates were incubated at the indicated temperatures for 2 days. Pictures of one representative isolate are shown for each genotype.

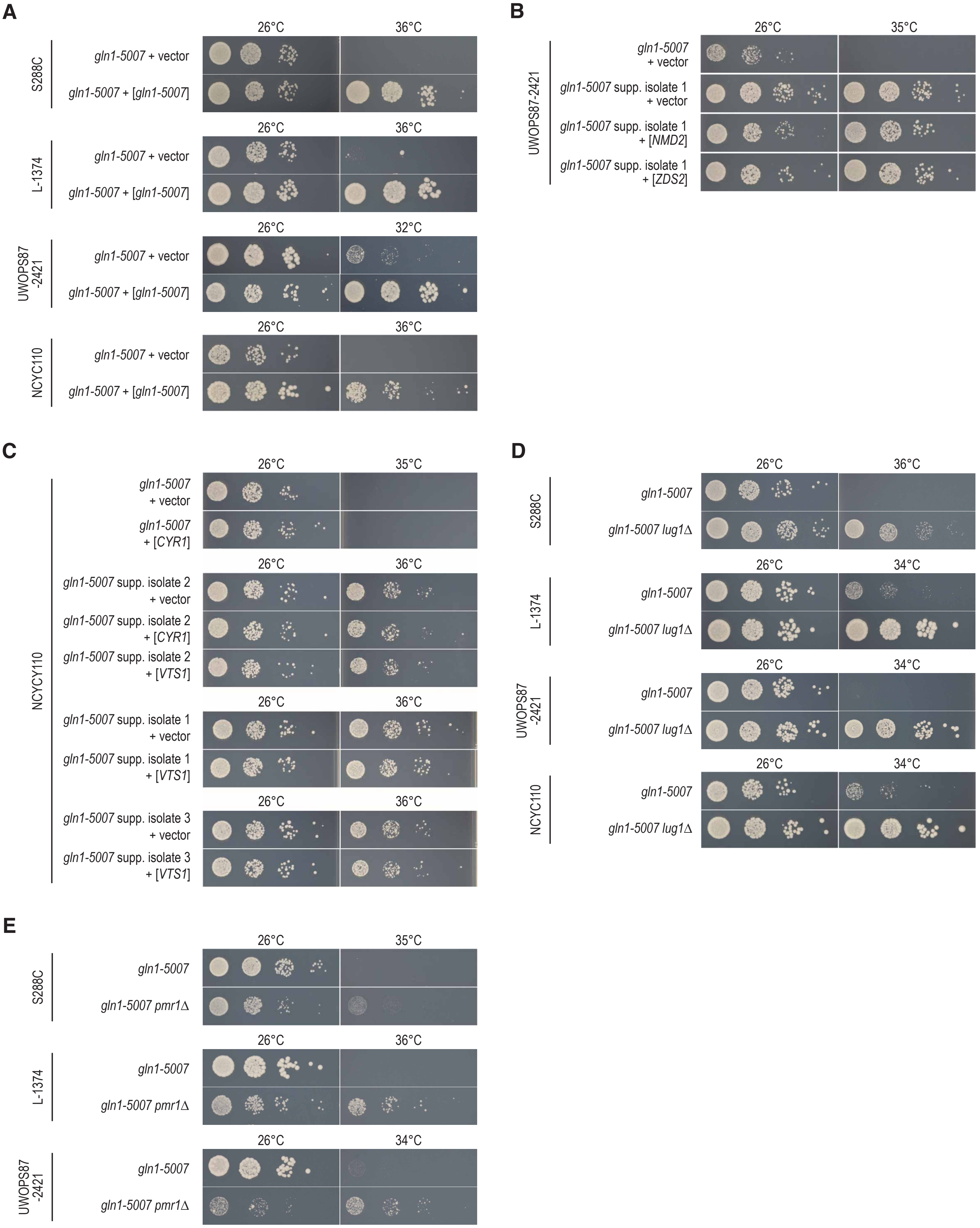


**Fig. S4. Validation of *GLN1* suppressors.** (**A**-**E**) Spot dilution assays of *gln1-5007* strains overexpressing *gln1-5007* (A), *NMD2* (B), *ZDS2* (B), *CYR1* (C), or *VTS1* (C), or deleted for *LUG1* (D) or *PMR1* (E). Cultures of 2-3 independent isolates of the indicated strains were grown until saturation, and a series of ten-fold dilutions was spotted on SD-Ura plates. Plates were incubated at the indicated temperatures for 2 days. Pictures of one representative isolate are shown for each genotype.

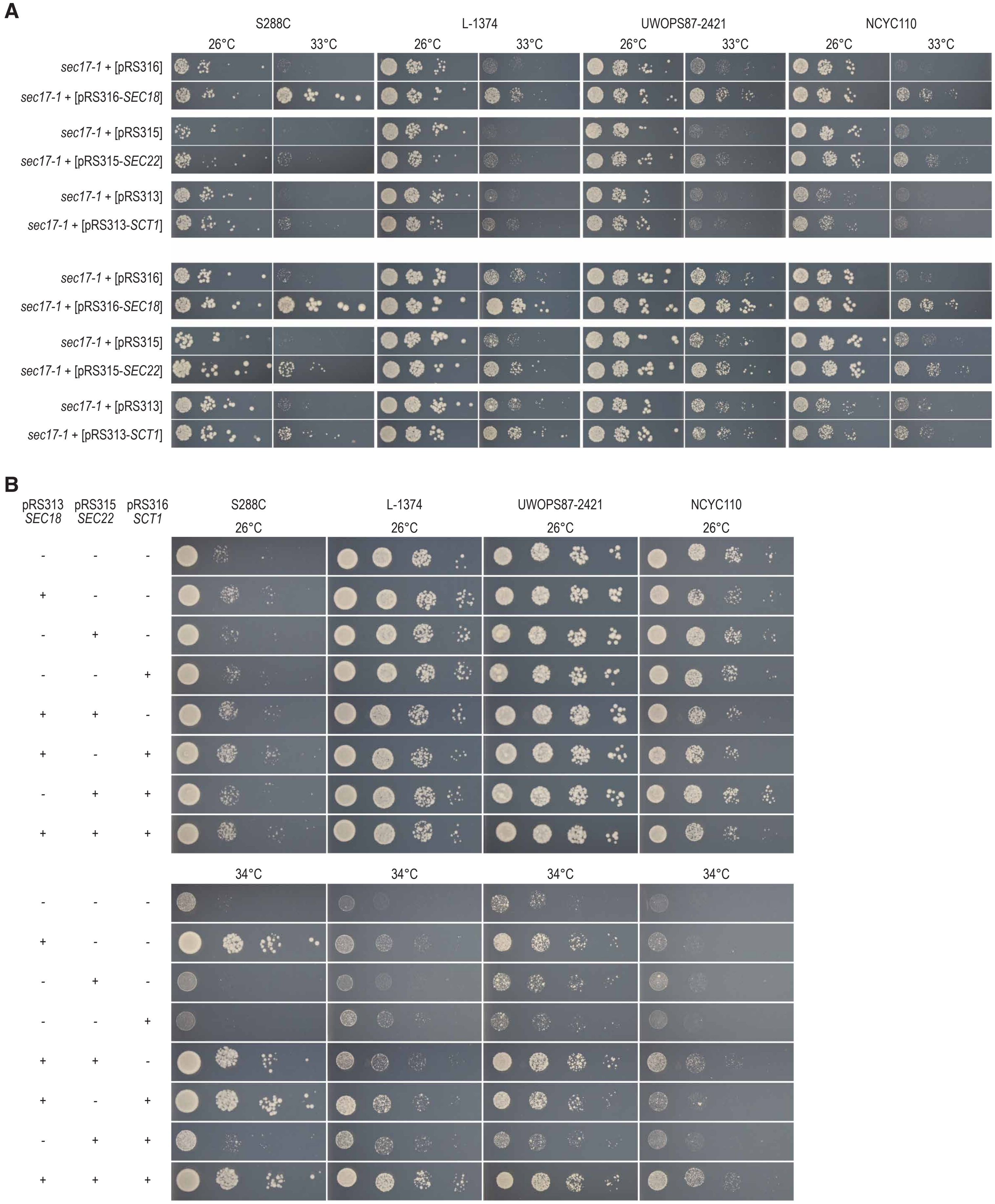


**Fig. S5. Multiple genes can contribute to the suppression phenotype.** (**A**) Validation of *sec17-1* suppressors using pRS-plasmids. *SEC18, SEC22*, or *SCT1* were cloned into pRS-plasmids and transformed into S288C, L-1374, UWOPS87-2421, and NCYC110 *sec17-1* strains. In each case, the *SEC18*, *SEC22*, and *SCT1* overexpression alleles matched the genetic background in which they were transformed, such that S288C was transformed with S288C alleles and L-1374 with L-1374 alleles, etc. Cultures of three indepdent transformants of the indicated strains were grown until saturation, and a series of ten-fold dilutions was spotted on SD-Ura (*SEC18* overexpression), SD-Leu (*SEC22* overexpression), or SD-His (*SCT1* overexpression). Plates were incubated at the indicated temperatures for 2 (top) or 3 (bottom) days. Pictures of one representative transformant are shown for each genotype. (**B**) Spot dilution assays as in (A), but using combinations of *SEC18*, *SEC22*, and/or *SCT1* plasmids. Plates were incubated at the indicated temperatures for 2 days. + = strains were transformed with the indicated plasmids. - = strains were transformed with the corresponding empty vectors.
